# *De novo* brain-computer interfacing deforms manifold of populational neural activity patterns in human cerebral cortex

**DOI:** 10.1101/2021.06.28.450263

**Authors:** Seitaro Iwama, Zhang Yichi, Junichi Ushiba

## Abstract

Human brains are capable of modulating innate activities to adapt to novel environmental stimuli; for sensorimotor cortices (SM1) this means acquisition of a rich repertoire of motor behaviors. We investigated the adaptability of human SM1 motor control by analyzing net neural population activity during the learning of brain-computer interface (BCI) operations. We found systematic interactions between the neural manifold of cortical population activities and BCI classifiers; the neural manifold was stretched by rescaling motor-related features of electroencephalograms with model-based fixed classifiers, but not with adaptive classifiers that were constantly recalibrated to user activity. Moreover, operation of a BCI based on a *de novo* classifier with a fixed decision boundary based on biologically unnatural features, deformed the neural manifold to be orthogonal to the boundary. These principles of neural adaptation at a macroscopic level may underlie the ability of humans to learn wide-ranging behavioral repertoires and adapt to novel environments.

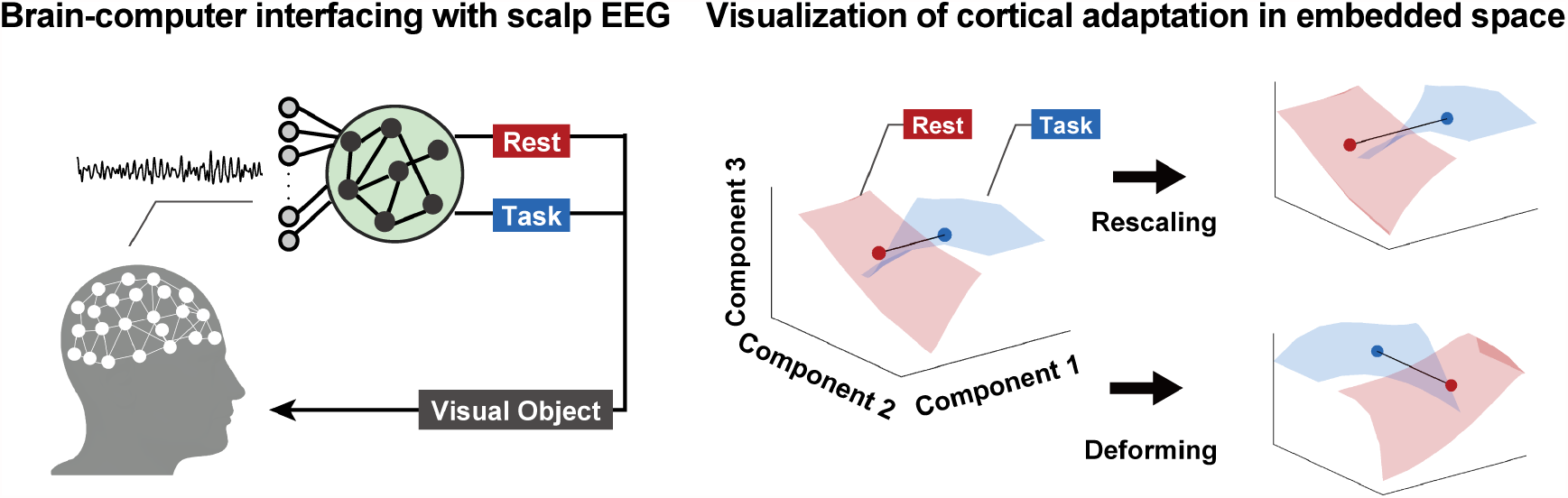

## 1 Introduction

Neural plasticity underlies behavioral adaptation to the external environment by changing properties of neural circuitries involved in, for example, dexterous motor behaviors, such as sports, musical performance, tool-use, or brain-computer interface (BCI) operations (Imamizu et al., 2000; Nudo et al., 1996; Quallo et al., 2009). The adaptation processes to achieve purposeful physical movement have been examined by electrophysiology, neuroimaging, and behavioral approaches (Karni et al., 1995; Kleim et al., 2004; Kording et al., 2007; Nudo et al., 1996; Shadmehr & Mussa-Ivaldi, 1994).

In sensorimotor studies leveraging neural activity recordings, local neuronal circuitries display repertoires of firing patterns that reliably represent ongoing behavior (Gallego et al., 2018; Shenoy & Kao, 2021). This representation of covariance structure has been referred to as the neural manifold, and intriguing findings suggest that the brain is capable of rapidly learning patterns of spike activities inside the manifold but not those outside of it (Sadtler et al., 2014). These constraints to learning, which are putatively due to the microscopic configuration of neurons, illustrate realistic behavior as well as the BCI control on which neural activities and behavioral consequences are directly mapped.

While a neural manifold describes the constraints on the ensemble of local neural activities in which hundreds of neurons are implicated, what is less investigated are the constraints on the macroscopic neural system. Because the brain exerts information processing via not only local circuitry but also the inter-regional coupling by which macroscopic neural populations selectively communicate (Bassett et al., 2015), those implicated in coherent communication might also be constrained similarly to the local circuitries (Fries, 2005, 2015).

To characterize the constraints on cortical population activities during adaptation, we used BCIs based on scalp electroencephalograms (EEG) with a variety of incorporated classifiers (Figure 1).

**Figure 1.**
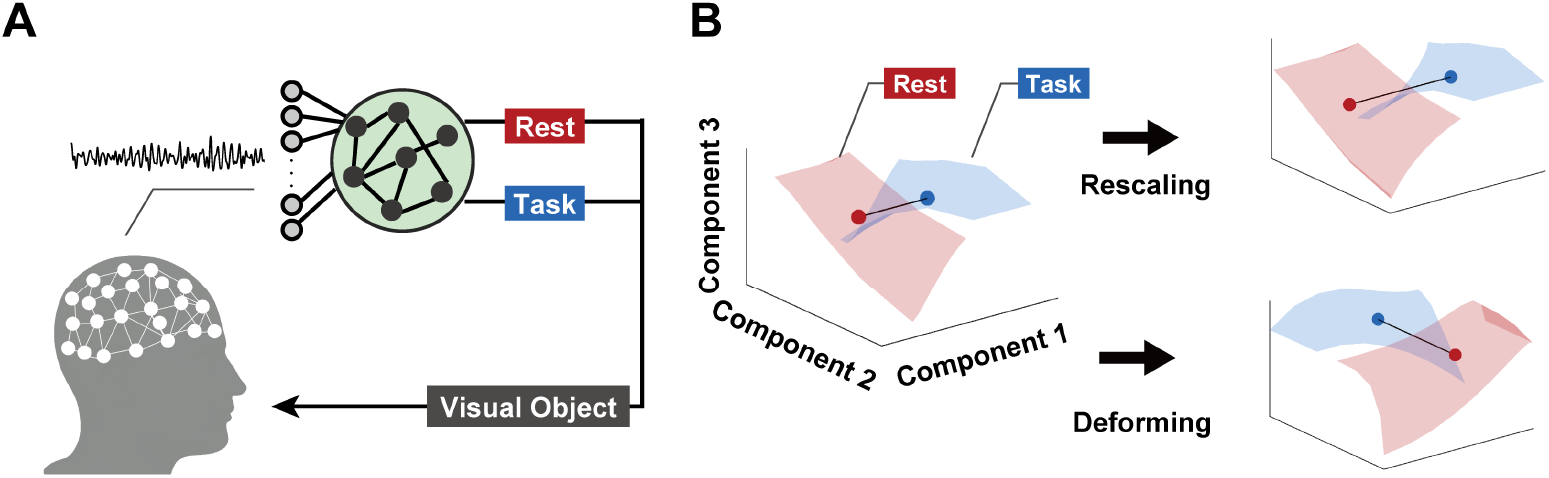
Conceptual illustration of neural adaptation process induced by brain-computer interfacing. A: Setup of a brain-computer interface. Online acquired scalp electroencephalograms were fed into a classifier to detect the presence/absence of attempted movement. Predicted brain state was shown to participants as movement of visual object on display. B: Conceptual visualization of cortical adaptation. Scaling adaptation reflects improvement in voluntary regulation of a specific component. If the centers of gravity determined from datapoints in two conditions are separated after brain-computer interfacing, it suggests the separability of two conditions is enhanced by adaptation. Deforming adaptation suggests that activity patterns are allocated to a specific brain state in order to adapt to the classifier. If the geometric relationships between two conditions are deformed with respect to a specific axis, it suggests the adaptation process progressed such that the two conditions are ’
sseparated along the axis.

Users attempted to move a virtual object using mental actions that modulated EEG signals. For each user, one of three classifiers determined the movement of objects based on a different set of rules. The model-based classifier required voluntary attenuation of sensorimotor rhythm (SMR) power derived from sensorimotor cortex (SM1). This fixed BCI operation rule is consistent with physiological findings, as the attenuation of SMR reflects SM1 excitability (Naros et al., 2019; Pfurtscheller & Lopes Da Silva, 1999; Takemi et al., 2013), as well as functional coupling among sensorimotor-related regions (Hayashi et al., 2020; Schulz et al., 2014; Tomassini et al., 2020; Wander et al., 2013). The adaptive classifier based on machine learning algorithms was configured based on recent whole-head EEG activity patterns to achieve maximum BCI controllability. This data-driven classifier configuration entails adaptive weighting on signaling features implicated in not only sensorimotor, but also attentional or cognitive functions in which the front-parietal network is implicated (Corsi et al., 2020). Lastly, the *de novo* classifier had a fixed configuration based on a biologically unnatural feature – desynchronized alpha oscillations derived from parietal regions. Due to the absence of prior knowledge to control this feature, users were encouraged to explore mental actions to control a visual object in the given BCI framework (Fujisawa et al., 2019).

As the decision boundaries between resting and motor attempts for each of the three classifiers (classifier plane) differed in their configurations, cortical adaptation processes were investigated by t-distributed stochastic neighbor embedding (t-SNE) algorithms, a nonlinear dimensionality reduction to visualize the distinct geometric changes of a whole-head EEG signal during operating/learning BCI tasks (Van Der Maaten & Hinton, 2008).

## 2 Results

### 2.1 Score acquisition during brain-computer interfacing

Twenty-one participants operated BCIs with one of three randomly allocated classifiers that provided scores contingent on BCI performance (Figure 2, Figure 2–supplement 1, 2). While BCI performance scores from the model-based and adaptive classifier generally increased over sessions, those for the *de novo* classifier did not. Statistical tests for coefficient of linear regression acquired from each participant revealed significant differences from zero for BCIs based on the model-based and adaptive classifiers (model-based: *p =* 0.0078, adaptive: *p =* 0.023, *de novo*: *p =* 0.055, Wilcoxon rank-sum test, FDR corrected). Note that direct comparison of the coefficients among classifiers is not possible because scores from each classifier were computed based on different classifiers.

**Figure 2.**
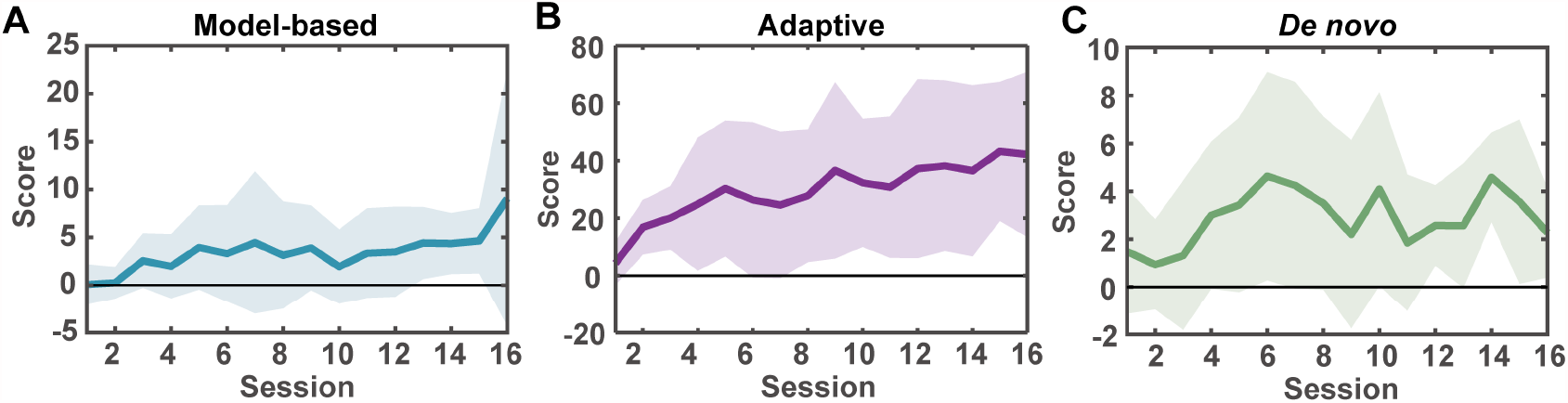
Temporal changes in acquired scores. Group results of performance scores from users of model-based (A), adaptive (B), and *de novo* (C) classifiers. Solid lines indicate mean values while shaded areas represent 1 standard deviation across participants.

### 2.2 Quantification of cortical adaptation process to classifier’s separating plane

To examine differences in cortical adaptation processes, we next investigated changes in whole-head EEG signals (Figure 3A). Using band-power features as a representation of brain state, all data acquired from a single experiment were subjected to the t-SNE algorithm to evaluate geometric relationships among two brain states (i.e., resting and attempted movement) and the classifier plane in the embedded space.

**Figure 3.**
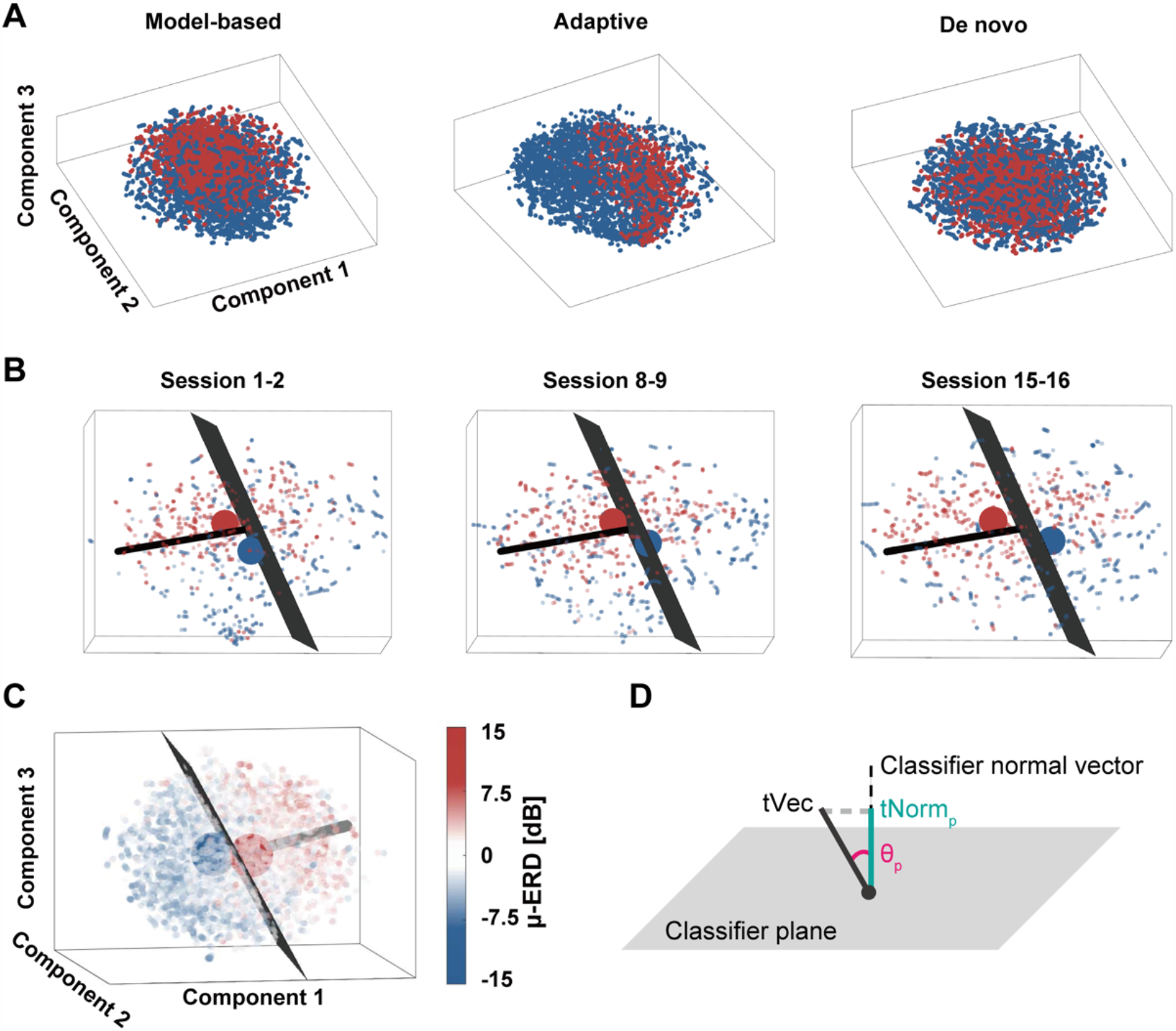
low dimensional visualization of EEG data by t-SNE. A: Examples of t-SNE-based visualization of datasets from a representative participant in each classifier. Each axis represents results of the t-SNE analysis, which generates three axes from input data. Blue points represent data from the Imagine period and red ones are those from the Rest period. B: Changes in geometric relationships between dataset and classifier plane. As training progressed, the geometric relationship of points from two brain states changed with respect to the classifier plane (black plane). The large points indicate the centers of gravity of points from each brain state. The black line orthogonal to the classifier plane is the classifier normal vector (see also Figure 3D) C: An example of t-SNE-based data visualization in embedded space (Model-based classifier user). Each datapoint is colored with its SMR-ERD value derived from the C3 electrode around the left sensorimotor cortex. The black plane represents the classifier plane (see also equation 2.9 for mathematical details). The large points indicate the centers of gravity of points from each brain state. The black line orthogonal to the classifier plane is the classifier normal vector (see also Figure 3D). D: The t-SNE-based quantification of the adaptation process with respect to the classifier plane. *tNorm*_*p*_ is defined as a component of *tVec* with respect to the classifier vector, while *θ*_*p*_ is defined as a subtended angle between *tVec* and the classifier vector.

An example of data from the model-based classifier BCI is shown in Figure 3B. As the participant performed the BCI operation, data during attempted movement (blue points) moved across the classifier plane, where the sign of relative SMR power flips (Figure 3C). In this case, the defined metrics *tNorm*_*p*_ and *θ*_*p*_ (Figure 3D) increased and decreased, respectively.

Figure 4A depicts changes in *tNorm*_*p*_ and *θ*_*p*_ between the first and last four sessions in the experiment. For participants trained with the model-based classifier, *tNorm*_*p*_ values significantly increased (*p* = 0.016, *d* = 0.71, two-tailed Wilcoxon signed-rank test) and the change was specific to participants who operated with model-based classifiers (Figure 4–supplement 1A, *p* = 0.81, 0.047). At the same time, *θ*_*p*_ values decreased significantly for participants trained with both the model-based and *de novo* classifiers (*p* = 0.016, *d =* 0.77, *p* = 0.016, *d* = 1.0, respectively, Figure 4–supplement 1B), but not with the adaptive classifiers (Figure. 4C; *p* = 0.58).

**Figure 4.**
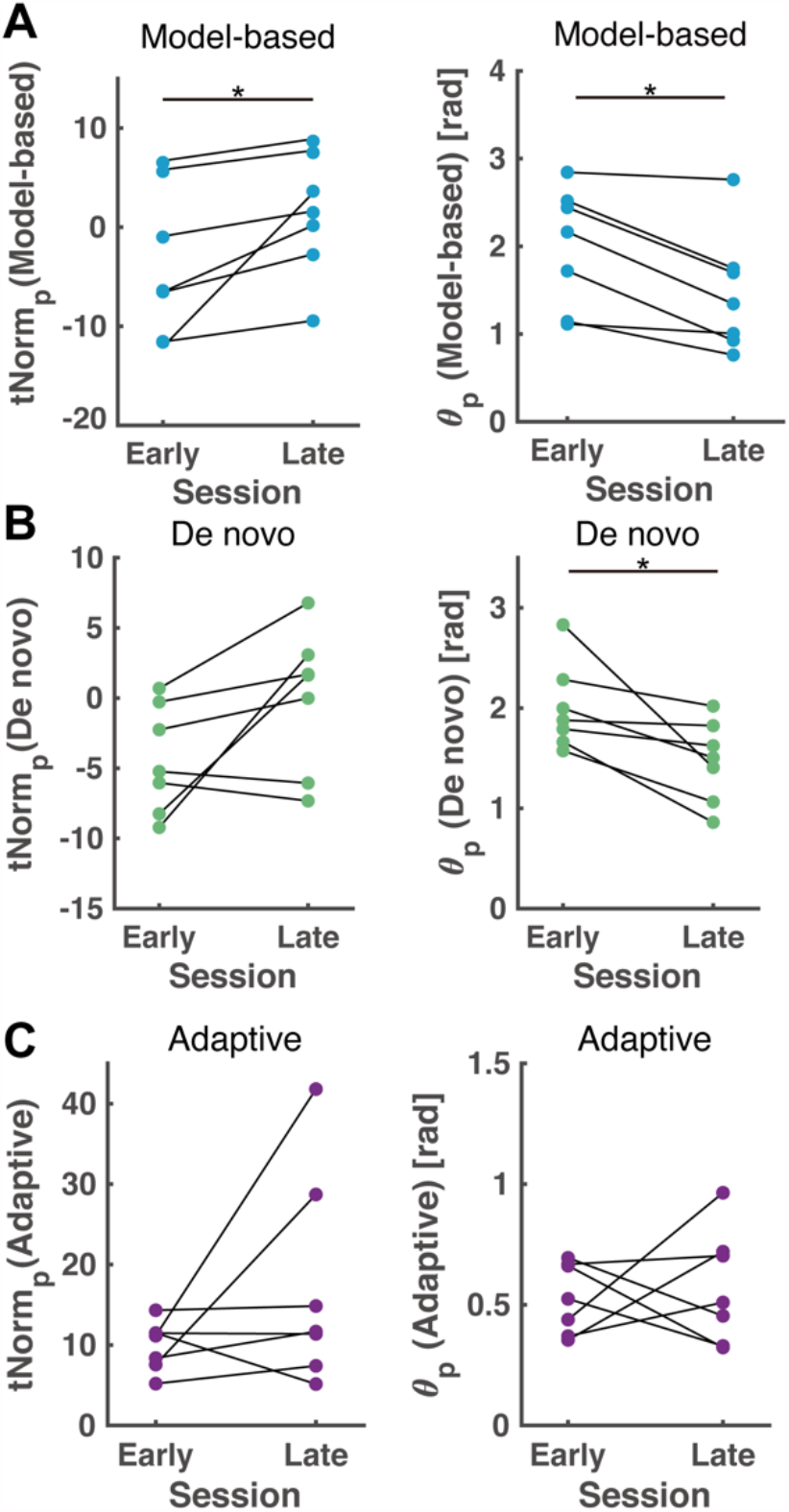
Quantitative comparison of cortical adaptation processes in embedded space. Changes over time in *tNorm*_*p*_ and *θ*_*p*_ for participants operating under the model-based classifier (A), the *de novo* classifier (B), and the adaptive classifier (C).

The identical evaluation was conducted for the *de novo* classifier plane. Figure 4B depicts changes in *tNorm*_*p*_ and *θ*_*p*_ against the *de novo* classifier. While no significant differences were confirmed for *tNorm*_*p*_ values over sessions (*p* = 0.078), *θ*_*p*_ values decreased significantly (*p* = 0.016, *d* = 1.3). Neither *tNorm*_*p*_ nor *θ*_*p*_ changed with respect to the other two classifiers (Figure 4–supplement 2, model-based: *p* = 0.047, 0.031, adaptive: *p* = 1, 0.58, *tNorm*_*p*_ and *θ*_*p*_, respectively).

As the classifier planes changed from one session to the next for the adaptive classifiers trained with the data from the previous sessions, each metric was calculated against the classifier plane determined with the dataset from the previous session. No significant differences were confirmed for comparison between the early and late period for the adaptive classifier (Figure 4C, supplement 3).

## 3 Discussion

In the present study, participants performed BCI operations with one of three classifiers: model-based, adaptive, or *de novo*. Each classifier elicited a different cortical adaptation process consistent with their characteristics. t-SNE analyses in embedded space revealed increases in *tNorm*_*p*_ for the model-based classifier, indicating rescaling of the neural manifold with respect to the axes orthogonal to the fixed decision boundary. Meanwhile, changes in population activities were not induced by the adaptive classifiers; decreases in *θ*_*p*_ indicated that the manifold was deformed, resulting in a reconfiguration orthogonal to its classifier plane by the *de novo* classifier that was based on biologically unnatural features.

### 3.1 Tuning classifiers to a brain induced by adaptive algorithm

Both the model-based and adaptive classifiers elicited short-term learning of the BCI operations as evidenced by the increases in performance scores; however, these two processes were distinct from one another. While model-based classifiers elicited changes in *tNorm*_*p*_ and *θ*_*p*_, the adaptive classifiers did not. Such a difference might be attributed to the design of the classifiers, as the mental actions that users were instructed to perform were identical. During BCI operations with a constant classifier plane, participants honed their abstract ability to control sensorimotor activity by minimizing error between the current classified result and their intended mental action; however, in the case of the adaptive classifiers, adaptation of users to the classifier was putatively limited due to the session-by-session recalibration.

Despite the absence of cortical adaptation to the classifier plane for users of the adaptive classifiers, performance scores did increase incrementally throughout the experiment. Accordingly, we can only posit that the adaptation of classifiers to users systematically progressed across sessions. It should be noted that implementing the adaptive algorithm might induce suboptimal results when the objective of the BCI operation is the induction of a specific neural activity, such as changes in excitability, activity patterns, or connectivity of targeted regions (Ramot et al., 2017; Ruddy et al., 2018; Shibata et al., 2011).

### 3.2 Cortical adaptation process during *de novo* brain-computer interfacing

Although significant increases in performance scores and *tNorm*_*p*_ were not confirmed for the *de novo* classifiers, cortical adaptations towards the classifier plane were partly observed, as evidenced by the decreases in *θ*_*p*_. The *de novo* task was defined as one that participants work on to improve their performance without any prior knowledge or strategy (Choi et al., 2020; Fujisawa et al., 2019; Radhakrishnan et al., 2008; Telgen et al., 2014). To achieve this during brain-computer interfacing, neurofeedback was provided via an illustrated tail. Because movement of a tail is not inherent for humans, participants were instructed to explore possible mental actions that might be suitable for operation. As such an exploratory strategy might require more extensive training than recalibrating the existing control configuration, performance scores did not tend to progress within a single-day experiment (Choi et al., 2020; Telgen et al., 2014).

The neural adaptation process was visualized via the t-SNE-based analysis. Deforming effects, that is rotational changes in the geometric relationship of two brain states towards the classifier plane, were confirmed in participants using the *de novo* classifier. However, the absence of a significant scaling effect suggested that the dissection of the two conditions (resting and motor imagery) did not systematically progress; this observation might reflect the reassociation of existing activity patterns to adapt to the BCI classifier by exploring a strategy to control the object. The result is consistent with the time course of the performance score and possible necessity of multi-day training to affect substantial behavioral improvement in *de novo* learning (Choi et al., 2020; Fujisawa et al., 2019). Although the flexibility of the human brain enabled partial adaptation to the *de novo* classifier planes, the adaptive classifier did not elicit brain-side adaptation. These findings collectively suggest that fixation of the classifier plane is an essential element for inducing neural plasticity via a brain-computer interaction based on macroscopic neural population activity.

## 4 Material and Methods

### 4.1 Participants

Twenty-one neurologically healthy adults (9 females, 12 males, mean age: 22.6 ± 3.23) who had never operated a BCI participated in this experiment. The appropriate sample size for this study was determined by an a-priori power analysis (α = 0.05, 1-β = 0.8, two-sided Wilcoxon signed-rank tests) focusing on the deforming effect induced by *de novo* BCI. The statistical package G*Power 3 (Faul et al, 2007) was used to estimate the sample size that shows large Cohen’s *d* = 0.90 reported in the previous EEG-based neurofeedback literatures (Hayashi et al., 2020; Soekadar et al., 2015). We calculated that 7 participants were needed.

All participants had normal or corrected-to-normal vision and were asked to provide written informed consent before participating in the experiment. This study was conducted according to the ethics of the Declaration of Helsinki. The experimental protocol was approved by the ethical committee of the Faculty of Science and Technology, Keio University (Approval Number: 2020-36, 31-23).

### 4.2 Experimental setup

Participants were seated on a comfortable chair in a quiet room. A display was placed about one meter in front of the chair to provide task instructions and visual feedback from BCIs.

EEG signals during the experiment were acquired with a 128-channel HydroCel Geodesic Sensor Net (HCGSN, EGI, Eugene, OR, USA.). The layout of channels followed the international 10-10 electrode positions shown in Figure 2–supplement 1A (Luu & Ferree, 2005). The reference channel was set to Cz. The impedance of all channels was maintained below 50 kΩ throughout the experiment. The EEG data were collected with a sampling rate of 1 kHz and transmitted via the Ethernet switch Gigabit Web Smart Switch (Black Box, Pennsylvania, USA) to EEG recording software Net Station 5.2 manufactured by EGI and MATLAB R2019a (The Mathworks, Inc, Massachusetts, USA).

### 4.3 Online processing of EEG signals

Analytical pipelines for online signal processing were implemented with a combination of MATLAB and Unity (Ver. 2019.2.4f1, Unity Technologies, USA). While the real-time status of brain states was determined from EEG signals processed by a custom MATLAB script, Unity presented the visual neurofeedback objects. EEG signals were processed with a 1651-point, minimum-phase, FIR 8-30 Hz bandpass temporal filter and then processed with one of the three types of BCI classifiers. Online processed EEG signals were used to classify brain-states as either presence or absence of attempted movement with one of the three types of classifiers: model-based, adaptive, or *de novo*. Each classifier was designed with different rules, and electrodes of interest were defined as shown in Figure 2– supplement 1A.

The model-based classifier was constructed based on those used in common SMR-based BCIs (Buch et al., 2008; Kraus et al., 2016). EEG signals around the left SM1 (i.e., channel C3) were only used to detect the attempted movement, because accumulated evidence suggests that event-related desynchronization of SMR (SMR-ERD) contralateral to the hand that attempted to move reflects the excitability of SM1 (Hummel et al., 2002; Naros et al., 2019; Takemi et al., 2013). In online processing, a large Laplacian filter was applied to EEG signals from channel C3 to extract sensorimotor activity (McFarland et al., 1997; Tsuchimoto et al., 2021). Subsequently, the band power of SMR (SMR-power; 8-13 Hz) was extracted by Fourier transform with a 1-s window and Hamming window function. The magnitude of SMR-ERD [dB] was computed from the obtained SMR-power with the following formula:

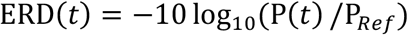

where P(*t*) denotes the power of interest, here the SMR-power, at time point *t*, and P_*Ref*_ denotes the reference power (Pfurtscheller & Lopes Da Silva, 1999). The reference power was calculated from the middle 3-s period of “Rest” time from the previous trial. The online calculated magnitude of SMR-ERD was then used as the index of neurofeedback for the model-based classifier. Movements of the illustrated hand in the display and performance scores were defined to be linearly related to the SMR-ERD value in the range of 0 to 10 dB.

The adaptive classifier was constructed using whole-head scalp EEG signals based on a common spatial pattern (CSP) algorithm and a support vector machine (SVM) (Blankertz et al., 2007). CSP components were extracted to maximize the separability of the two conditions Rest and Imagine, and were quickly trained at the end of each session to adapt to the current activity patterns of users. Specifically, the CSP was generated from the spatial covariance matrices of all EEG electrodes to find linear combinations of electrodes to form spatial filters that maximized the variance difference between the two conditions. The corresponding variances of spatially filtered EEG data were then divided into time windows and log-transformed to transform their distribution into a normal distribution. The SVM classifier was constructed to perform a binary classification of the two conditions. The posterior probability for a data point classified as presence of motor attempt was used as an index for neurofeedback; the index for the adaptive classifier was defined to be linearly related to the posterior possibility in the range of 50% to 100%. Note that the rules for object movement were identical to those of the model-based classifier, only the feedback was different.

Lastly, the *de novo* classifier had a fixed classifier plane as did the model-based classifier; however, its characteristics were biologically unnatural; the *de novo* classifier was based on EEG signals around the parietal region (i.e., channel Cz) that are associated with attentional features but not with sensorimotor activity (Benedek et al., 2014; Misselhorn et al., 2019). During the BCI task, users attempted to move their body or a visual object on the display; however, spectral power in the alpha-band (8-13 Hz) was increased by the motor attempt of moving the feet or by internal attention at the targeted channel (Benedek et al., 2014; Pfurtscheller et al., 2006). Such intrinsic responses did not contribute to the BCI operation, as the *de novo* classifier discriminated motor attempts with ERD values (i.e., power attenuation) in the alpha-band from the Cz channel, calculated with the procedure identical to that from channel C3 in the model-based classifier. Online computed ERD magnitude was exploited to decode the absence/presence of attempted movement and index for neurofeedback. Note that the rules for object movement and for obtaining scores were identical to those in the other two types of classifiers.

### 4.4 Experimental procedure

Participants underwent 16 BCI operation sessions, each consisting of 20 trials. All experimental procedures were conducted within 2 hours to guarantee the reversibility of any potentially unnaturally induced neural plasticity and to investigate the initial phase of learning to operate the BCIs. After every two sessions, participants were given a break of up to 5 min. Participants were randomly allocated to one of the three classifiers without knowledge of their existence or configuration and used the allocated type of classifier throughout the entire experiment.

A trial began with a 5-s “Rest” period. This was followed by a 5-s “Imagine” and a 3-s “Break” period (Figure 2 – supplement 1B). During the “Rest” period, participants were instructed to relax without having any specific thoughts and with opened eyes. In the “Imagine” period, participants were instructed by the experimenter to perform motor imagery tasks based on the allocated classifier. Participants with the model-based and adaptive classifiers were instructed to imagine extending the right-hand throughout the experiment, matching the imagined movement with the object on display. Participants with the *de novo* classifier were instructed to imagine moving a tail, also matching the movement of the object on display. As tail moving is not intuitive for humans, at the beginning of the session, participants were encouraged to exploratorily find a strategy that achieved the best control of the BCI. The strategy adopted in each session was freely determined by each participant, but they were instructed to try to use the same strategy throughout one session to acquire sufficient data during a specific strategy.

The performance of each trial was quantified by a score, which participants were encouraged to make as high as possible. Scores were determined by the predicted presence/absence of the attempted movement. Both the absence of attempted movements during “Rest” periods and the presence of attempted movements during “Imagine” periods resulted in higher scores, while scores were reduced if movements contrary these predictions were detected. The changing rates of these scores were pertinent to the metrics used for feedback by each classifier and were regulated linearly to fit the score range from minus one hundred to plus one hundred. After each session, participants were asked to verbally describe the strategies they had adopted.

For the adaptive classifier, the CSP-SVM model was re-trained with “Rest” and “Imagine” period data from the previous session to use in the next session. Note that the first session of the adaptive classifier task was identical to that of the model-based one, so as to collect a dataset for constructing the adaptive classifier. Detailed procedures for classifier training are described in the supplementary materials.

### 4.5 Evaluation of BCI performance

Online-calculated scores were subjected to linear regression analysis (Gruzelier, 2014; Kober et al., 2018; Witte et al., 2018). The score obtained during a given session was used as a dependent variable and session number was used as a predictor valuable. If scores increased during the experiment, the regression coefficient for the predictor valuable was positive. To test whether the obtained regression coefficients were significantly different from zero, they were subjected to a group-by-group Wilcoxon rank-sum test with a false discovery rate correction (Benjamini-Hochberg method; Benjamini & Hochberg, 1995).

### 4.6 Offline EEG preprocession

The recorded EEG signals were first preprocessed with EEGLAB (Delorme & Makeig, 2004) to reject artifacts and enhance the computational efficiency via downsampling (Bigdely-Shamlo et al., 2015). The raw EEG data were filtered with a zero-phase 1-45 Hz FIR bandpass filter and down sampled to 100 Hz. Channels classified as “Bad” by the EEGLAB plugin: Christian’s clean_rawdata (Bigdely-Shamlo et al., 2015) were removed from further analysis. The removed channels were interpolated spherically to minimize a potential bias when re-referencing the electrodes to a common average reference. Subsequently, large-amplitude artifacts caused by blinking or head displacement were removed via Artifact Subspace Reconstruction (Kothe & Makeig, 2013). The electrodes were then re-referenced to the common average reference to extract activity specific to the electrodes (McFarland et al., 1997).

The continuous EEG data were then segmented into trials to evaluate the middle 8-s periods of the online BCI training trials (i.e., the last 4 s of the “Rest” period and the first 4 s of the “Imagine” period). To obtain the independent EEG components of the segmented dataset, we used adaptive mixture independent component analysis (AMICA; Palmer et al., 2011). Finally, an automatic artifact rejection was applied using ICLabel that distinguished brain–originated EEG components from artifacts induced by eye, muscle, heart, line noise, and channel noises (Pion-Tonachini et al., 2019).

To investigate cortical adaptation processes during brain-computer interfacing, the band-power features were used as a raw-vector that represents instantaneous overall brain state. Computed band-power from each EEG channel was subdivided into five functionally distinct frequency bands (Delta: 1-4 Hz, Theta: 4-8 Hz, Alpha: 8-13 Hz, Beta: 13-31 Hz, Gamma: 31-45 Hz; Hayashi et al., 2019). The averaged band-power was log-transformed and normalized to the z-score in a trial-by-trial manner to cancel base-line drifting. Thereby, the original number of dimensions of the feature vector *D* was *D* = 129 × 5 = 645. The calculated band-power signals in the alpha-band also were subjected to cortical source estimation (See also supplementary materials).

### 4.7 Feature extraction of EEG-dataset using t-SNE algorithm

The preprocessed EEG dataset (645×11520 matrix) was subjected to a subject-by-subject t-SNE analysis, which converted the pairwise distances between data points in the original feature space to conditional probabilities (Van Der Maaten & Hinton, 2008). Mathematical details of t-SNE are further described in the Supplementary Materials and briefly here. First, the conditional probability that the data points *x*_*i*_ and *x*_*j*_ are neighbors was calculated from the pairwise distances of input data. Then, to maintain the probabilities in the original feature space in the embedded space, the Kullback-Leibler divergence representing the distance between the conditional probability in the original and embedded space was minimized via optimization. In this study, the number of dimensions of EEG features was reduced to three via a Barnes-Hut variation of t-SNE (Van Der Maaten et al., 2014) to speed up the computation. Perplexity, a hyperparameter of the t-SNE algorithm, was set to 20, which was determined empirically with a parameter search of past EEG data for best separation between the “Rest” and “Imagine” periods. The hyperparameter was fixed across participants throughout the study after the determination. After applying t-SNE, the dimensionality-reduced datasets were subjected to visualization and a similarity analysis.

### 4.8 t-SNE-based dimensionality reduction and quantitative analysis in embedded space

Feature extraction using dimensionality reduction is popularly conducted for high-dimensional neural data across modalities (Cunningham & Yu, 2014; Lord et al., 2019). The t-SNE algorithm we adopted has advantages for geometric evaluation, as it preserves original distances in the embedded space. Here, to conduct quantitative analysis beyond its general purpose for data visualization, we employed metrics that do not violate the assumption of the t-SNE algorithm. Feature extraction techniques such as ICA, principal component analysis, or factor analysis display weighted maps of extracted components so that they can be applied to newly acquired data, whereas the t-SNE algorithm does not. To enable interpretation of the dataset with reduced components, in the present study, entire datasets from individual participants were subjected to t-SNE analysis at once.

Because t-SNE unfolds the nonlinear structure of a given dataset, the linear distance in the embedded space can be interpreted as an approximation of geometric distance in the original space. It illustrates how different one brain activity pattern is from another. It should be noted however that to properly interpret the results (1) distance scales in the embedded space were rearranged and were variable across iterations of t-SNE, (2) distance scales in different clusters might have differed, (3) direct comparisons of distances between clusters were not acceptable because distances within two clusters were arbitrary. To deal with the above concerns, two approaches were adopted: (1) data points were bridged to prevent the formation of multiple clusters, and (2) statistical distances, namely Hotelling’s t-squared statistical values, were used instead of Euclidean metrics.

Because distances between nearby points are well preserved in embedded space, the distance scale of distant points were kept similar for enough data points, which acts as a bridge and prevents the formation of sparse multiple clusters. We also adopted the concept of “short-circuiting” (Lee & Verleysen, 2005) by constructing the feature vectors with overlapped time-windows so that points were smoothly connected. Thus, distances from point to point shared the same scale across all points (i.e., only one cluster was generated in embedded space).

Hotelling’s t-squared statistic was adopted as the distance metrics between two group of points (Hotelling, 1992). Assume *x* and *y* are two groups of points lying in a *p*-dimensional space, *n*_*x*_ and *n*_*y*_ are the numbers of points, 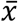 and 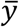 are the sample means, and 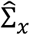 and 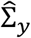 are the respective sample covariance matrices. The Hotelling’s t-squared statistic was calculated as:

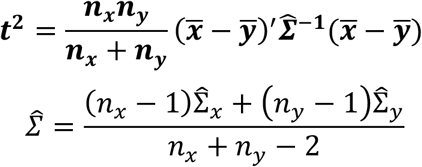

Hotelling’s t-squared statistic is suitable for measurements of statistical distance in the t-SNE-embedded space, as they were invariant to the distance scale. The distribution of *t*^2^ follows an *F*-distribution:

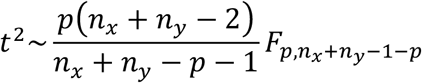

To normalize the distribution, the square root of *t*^2^ was defined as *tNorm* and was used as the distance measurement in subsequent analyses:

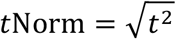

The vector representing the directional relationship between two classes was defined as *tVec*:

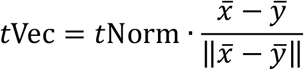

Data points were divided into two classes: “Rest” and “Imagine” according to their relative times in the trials. *tNorm* and *tVec* were calculated for these two conditions.

### 4.9 Classifier plane

To investigate the influence of BCI classifiers on the cortical adaptation in the t-SNE-embedded space, the classifier plane and classifier normal vector were linearly projected into the embedded space (See Figure 3C). The classifier vector *V* = [*v*_1_, *v*_2_, *v*_3_]^*T*^ was calculated as follows, where T denotes a matrix transpose.

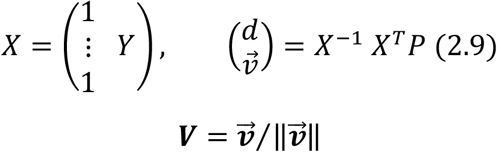

Then, the equation of the classifier plane is given as follows.

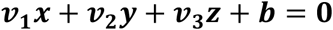

assuming *Y* ∈ ℝ^*N*×3^ are the points in the 3D embedded space, *P* ∈ ℝ^*N*×*n*^ are the original features referred to by the classifier, where *N* is the number of points, and n is the number of features. ***b*** is the intercept corresponding to the decision boundary of the classifiers.

As is shown in Figure 3D, *tVec* could be projected to the classifier vector to evaluate its positional relationship against the classifier. The lengths of projection on the classifier vector (*t*Norm_p_) and the angles between *tVec* and the classifier vector and (*θ*_*p*_) were calculated against that of the model-based classifier to directly compare the adaptation processes across classifiers as follows:

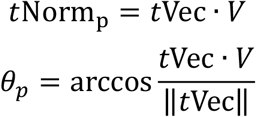

which represent the strength of scaling and deforming against the classifier plane, respectively.

### 4.10 EEG-similarity analysis

Geometry-based analysis was conducted in the embedded space, as positional relationships of the points reflected the similarities in the original space. The transition process from one brain condition to another (i.e., absence to presence of attempted movement) was assessed by the spatial arrangement and separability of points from the “Rest” and “Imagine” periods in the t-SNE dimension. Emergence of the two temporal phenomena were defined as follows:

- Scaling: The separability of the two conditions (Rest and Imagine) increases with respect to a fixed axis. Scaling is interpreted as the enhancement of specific cortical activity patterns.
- Deforming: The relationship of positions in the two conditions changes direction. Deforming is interpreted as an alteration of a cortical activity pattern that is adopted.

To quantify the two distinct adaptation process, the following metrics were defined. Scaling and deforming between the *i*^th^ and *j*^th^ sessions were quantified by *tNorm*_*p*_ and *θ*_*p*_.

- Scaling: Δ*t*Norm_*p*_ = *t*Norm_*p*_(*i*) − *t*Norm_*p*_(*j*)
- Deforming: Δ*θ*_*p*_ = *θ*_*p*_(*i*) − *θ*_*p*_(*j*)

If adaptation progresses toward the targeted neural activity patterns required to control BCIs, the *t*Norm_p_ values should be larger while those of *θ*_*p*_ should be smaller. Thus, the calculated values were subjected to the Wilcoxon sign-rank test to compare the differences between the first and last four sessions (early and late period, respectively). For adaptive classifiers, as the classifier plane was obtained from the 2nd session, the comparison was conducted from the 2nd session across all statistical tests. We then corrected the alpha-level with a Bonferroni correction.

## Acknowledgements

This study was supported by the Keio Institute of Pure and Applied Sciences (KiPAS) research program, JSPS KAKENHI Grant Number 20H05923 (to J.U.) and JST, CREST Grant Number JPMJCR17A3 (to J.U.) including the AIP challenge program, Japan. We thank Yumiko Kakubari, Shoko Tonomoto and Aya Kamiya for their technical supports.

## Figure Legends

**Figure 2 Supplement 1.**
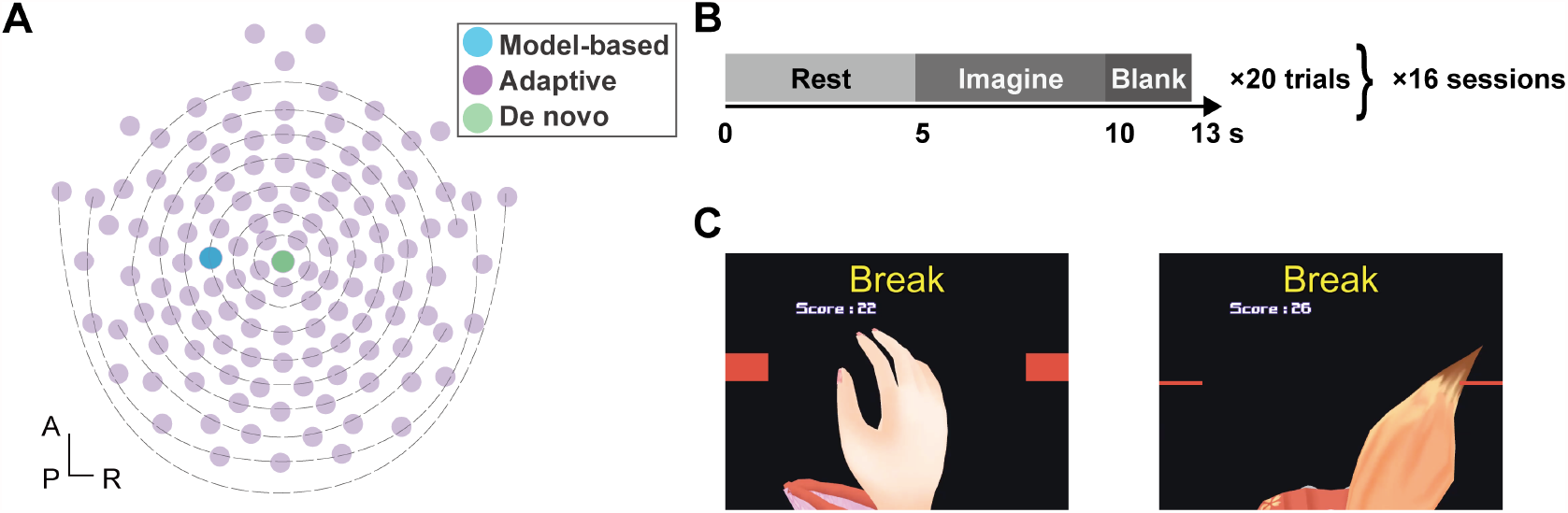
A: Electrode locations. The three classifiers used in the study had different channels of interest. The model-based classifier used only channel C3 indicated in blue around the left sensorimotor cortex. The adaptive classifier used whole-head EEG channels (purple) to construct a common spatial pattern. The *de novo* classifier used only the Cz channel, shown here in green. B: Experimental protocol and time course of a trial C: Visual feedback object. For the model-based or adaptive classifiers, an illustration of a hand was shown <u>**t**</u>hat matched the attempted movements of the users while an illustration of a tail was used in the *de novo* task to encourage users to acquire novel mental actions that enhanced controllability of the BCI.

**Figure 2 Supplement 2.**
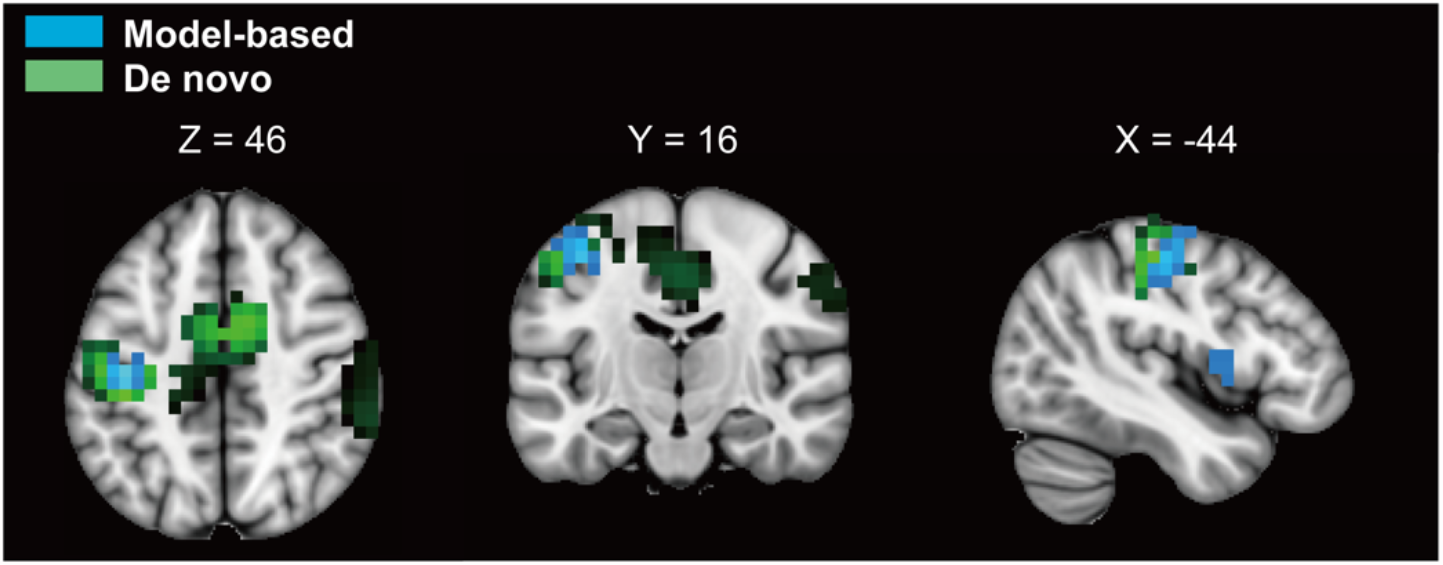
Results of source estimation analysis from representative participants. The colored regions indicate voxels where activities were significantly different during Rest and Imagine periods (*p* < 0.05 unc.). Areas colored with blue and green indicate those for model-based and de novo classifiers, respectively. While significant voxels were localized around the contralateral hemisphere of the imagined hand for the model-based classifier, those for the *de novo* classifier were located bilaterally, including in the pre/post central gyrus and supplementary motor area (peak voxel was in the postcentral gyrus, [MNI coordinates: -40, -25, 45]). Note that a representative source estimation for the adaptive classifier is not shown due to variable activity patterns among participants. sLoreta analyses of statistical non-parametric mapping for estimated cortical sources of band power in the alpha band (8-13 Hz). Areas colored with blue and green indicate those from model-based and *de novo* classifiers, respectively. Masks superimposed on a standard brain template were visualized by MRIcroGL (https://www.mccauslandcenter.sc.edu/mricrogl/home).

**Figure 4 Supplement 1.**
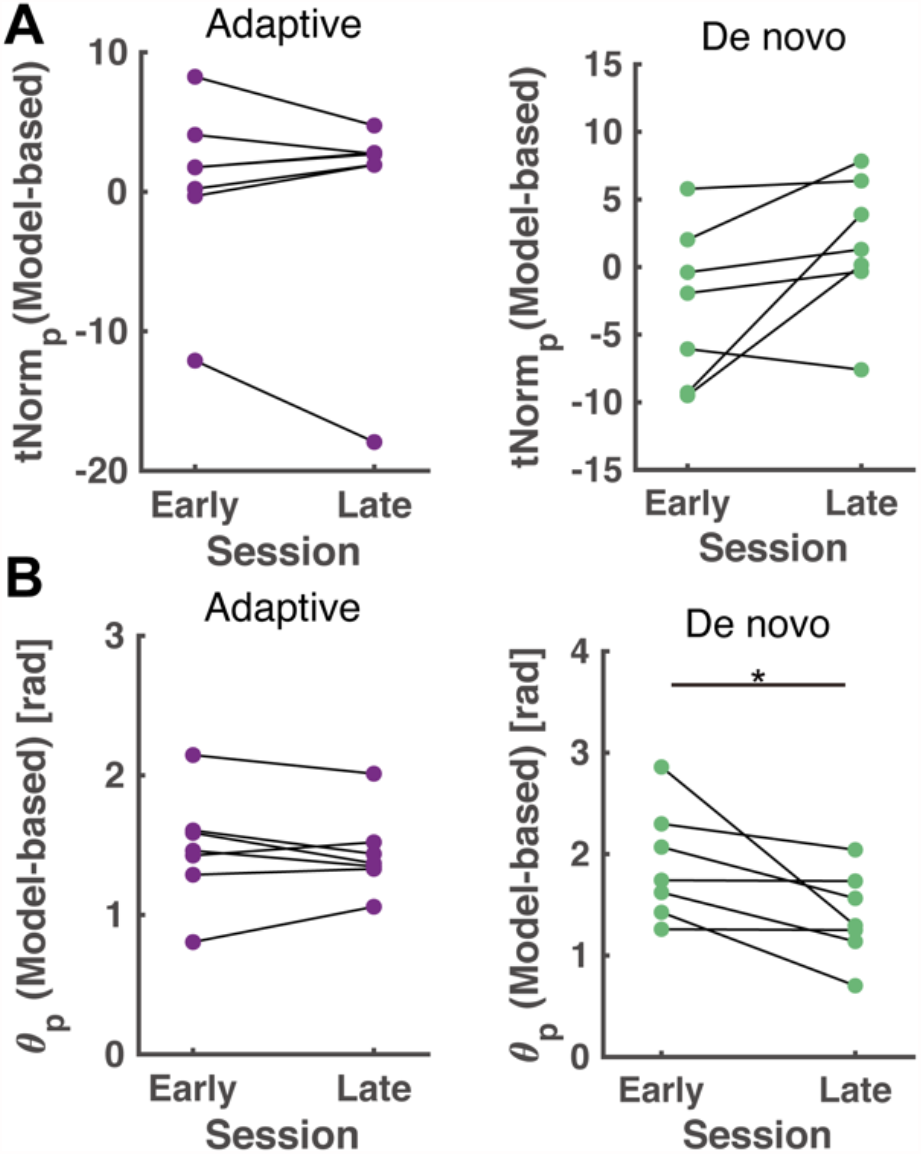
Quantitative comparisons of cortical adaptation processes to the model-based classifier plane Changes over time in *tNorm*_*p*_ (A) and *θ*_*p*_ (B) of the model-based classifier for participants operating under the adaptive (left, purple) or *de novo* (right, green) classifiers.

**Figure 4 Supplement 2.**
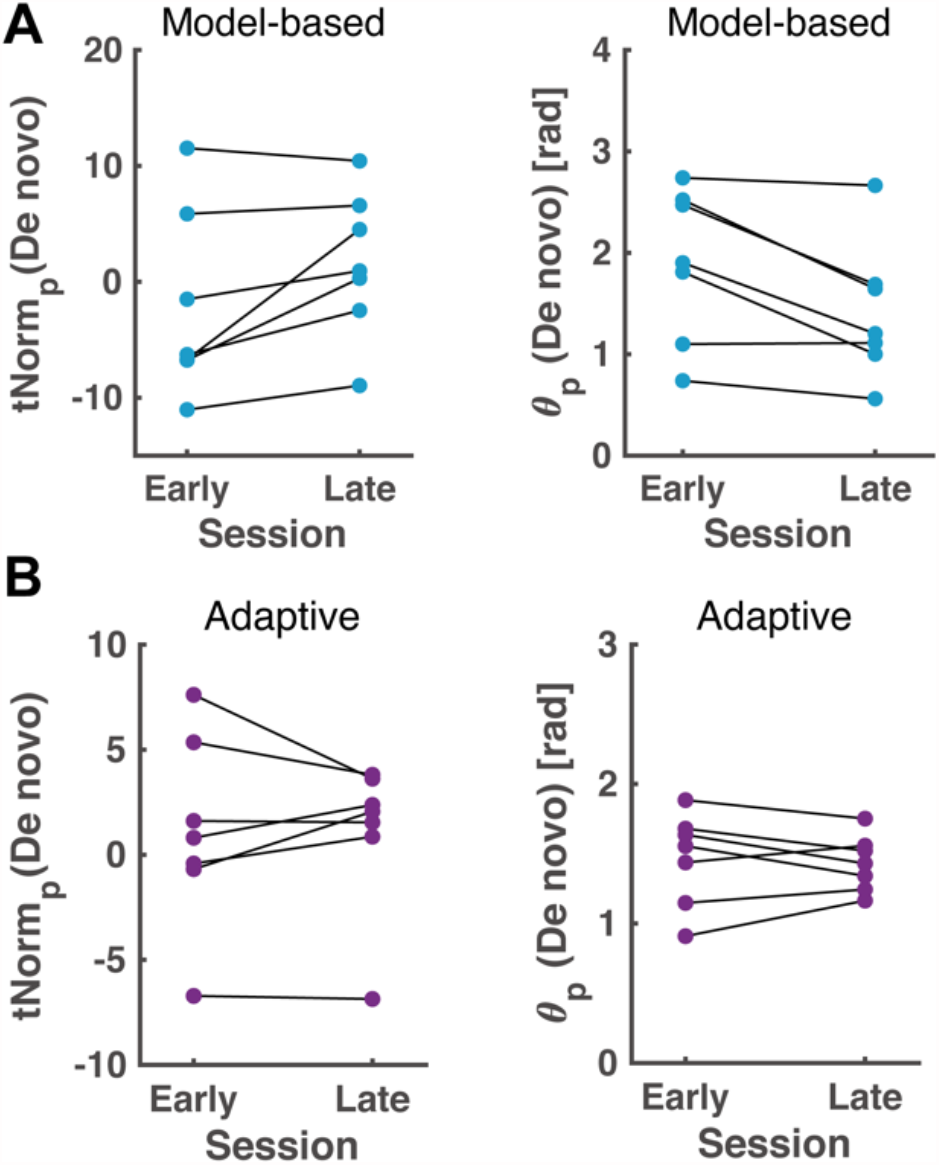
Quantitative comparisons of cortical adaptation processes to the de novo classifier plane. Changes over time in *tNorm*_*p*_ (left) and θ_p_ (right) of the *de novo* classifier for participants operating under the model-based (A, blue) and adaptive (B, purple) classifiers.

**Figure 4 Supplement 3.**
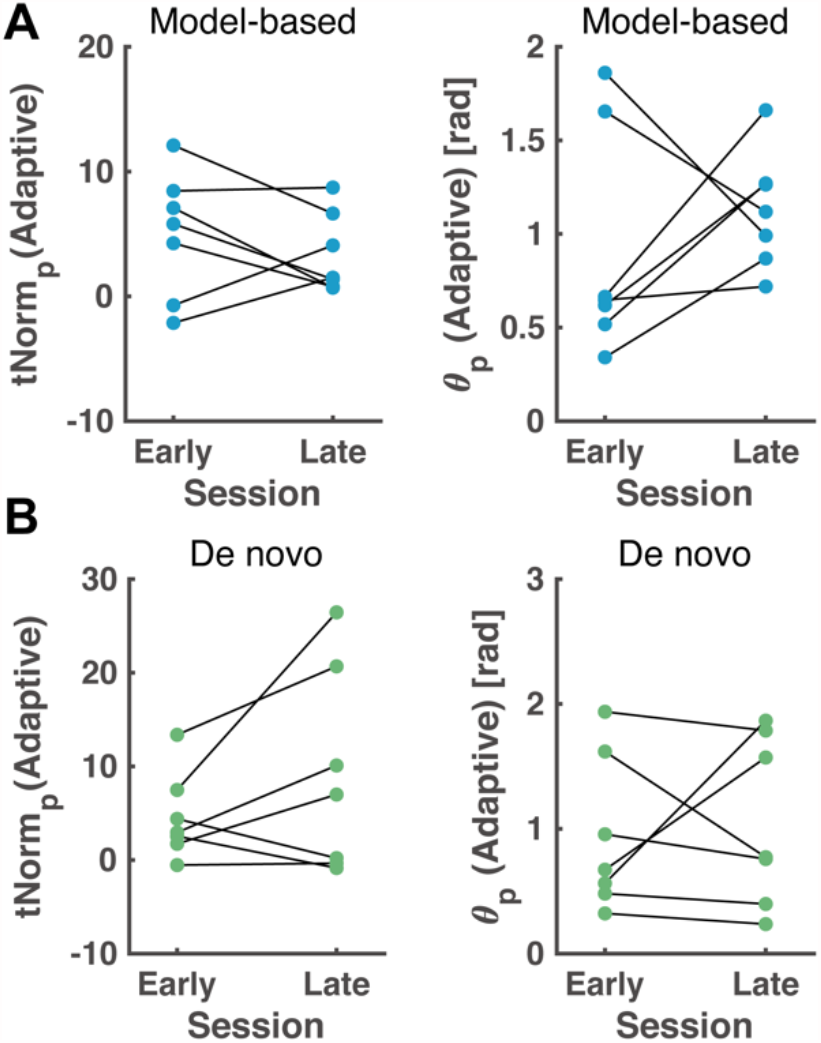
Quantitative comparisons of cortical adaptation processes to the adaptive classifier plane. Changes over time in *tNorm*_*p*_ (left) *and θ*_*p*_ (right) of the adaptive classifier for participants operating under the model-based (A, blue) and *de novo* (B, green) classifiers.

## Supplementary Material

### Detailed procedure for construction of adaptive classifier

In the experiments with the adaptive classifier, six spatial filters were generated via the common spatial pattern algorithm during the intervals between sessions. Filters that maximized variance differences were generated via the CSP algorithm and applied to the online EEG signals during the subsequent session. The log-transformed variances of the six-channel, spatial-filtered data from the previous 1-s signals were calculated and classified with a linear SVM classifier.

Cross validation results of the adaptive decoder are shown in Figure S1A. Data from a single session were used for training and data from the remaining sessions were used for testing. Performance, evaluated from the coefficients of linear regression, did not show systematic improvement at the group level (*p* = 0.078, Wilcoxon sign-rank test). Figure S1 B shows the results of a cross validation test using a single session for training and another for testing, suggesting there was an increase in accuracy during the later period, which was confirmed by temporal changes in the acquired score (Figure 2B).

**Figure S1.**
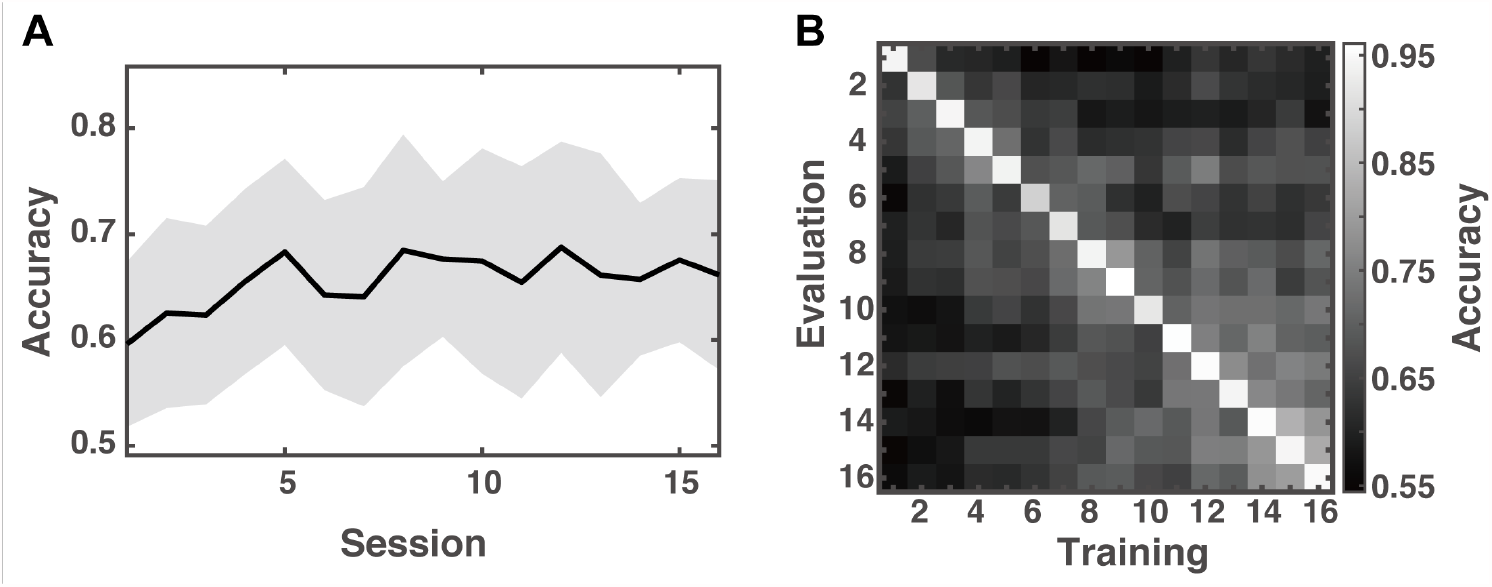
Temporal changes in cross validation performance of adaptive decoder

### Cortical source estimation analysis for EEG signals

For the data from participants who operated under the model-based or *de novo* decoders, band-power in the alpha-band was subjected to sLORETA analysis for cortical source estimation (Pascual-Marqui, 2002). Because the motivation for conducting the source analysis was to test whether the targeted region of the classifier was successfully activated during the late period of BCI training, averaged data from across the last four sessions were subjected to a non-parametric permutation test (Nichols & Holmes, 2002).

### Mathematical description of t-SNE algorithm

In the original manuscript describing the t-SNE algorithm (Van Der Maaten & Hinton, 2008), the datapoints are described as ***x*** = (*x*_1_ *x*_2_, … *x*_*n*_), where *x*_***i***_ is a vector with the arbitrary number of features. Assume that *x*_***i***_ and *x*_***j***_ are two data points, *d*_*ij*_ is the distance between the two points and *p*_*j*|*i*_ is the probability that *x*_***i***_. and *x*_***j***_ are neighbors. The probabilities follow a Gaussian distribution, described by:

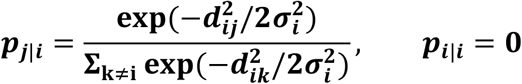

where *σ*_*i*_ is determined by the parameter *perplexity*, the value of which was calculated with *H*:

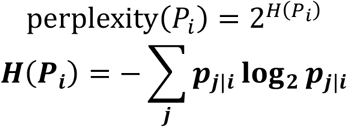

The value of *σ*_*i*_ is adjusted in a binary search method so that *perplexity* matches a value determined by the user. According to Van Der Maaten & Hinton (2008), perplexity is a smooth measure of the effective number of neighbors. The joint probability *p*_*ij*_ is defined by symmetrizing the conditional probabilities:

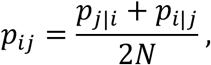

where *N* is the number of data points. The pairwise distances between points in low-dimensional embedding were converted to possibilities that follow a Student’s t-distribution with one degree of freedom. Assume that *y*_*i*_ and *y*_*j*_ are two data points in the embedded space and joint possibilities *q*_*ij*_ are defined as:

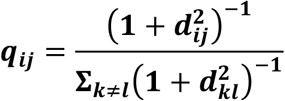

The Kullback-Liebler (KL) divergence between joint possibility distribution *P* in the original space and *Q* in embedded space was then calculated with:

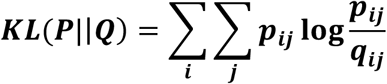

The KL divergence was minimized via a gradient descent method by adjusting the positions of points in the embedded space.

